# Di(2-ethylhexyl) adipate (DEHA), A NEW HOPE! A sustainable and promising process for the plasticisers industry

**DOI:** 10.1101/2020.10.26.355289

**Authors:** Federico Acciaretti, Andrea Pasquale

## Abstract

Plasticisers are commonly incorporated in plastic materials in order to improve their physico-chemicals properties. In particular, Poly-(vinyl chloride) (PVC) is a polymer which has excellent plasticiser compatibility characteristics. The demand for plasticized-PVC is steadily increasing and its synthesis need to be more sustainable, considering the interest in developing a circular economy in the next years. In order to achieve these goals, a bio-based process to synthesize di(2-ethylhexyl) adipate (DEHA), a widely used plasticiser, could be an interesting approach. The most important starting material for the process is adipic acid, but its synthesis from petrochemical sources is not sustainable. An alternative is using waste materials as substrates for fermentation in a totally green process. Among many strategies, the reverse adipate degradation pathway (RADP) in *E. coli* seems to be the most interesting one, considering the highest titer of 68 g/L and the yield of 93.1%. The next step is the enzyme-catalysed esterification of adipic acid and 2-ethylhexanol to produce DEHA, using an immobilized lipase from *Candida antarctica*. Applying a solvent-free system under vacuum condition is convenient as it guarantees a conversion to DEHA of 100 mol%.

## 1. Introduction

Plasticisers are a class of chemicals compounds added to a specific material to provide new and particular physico-chemicals characteristics. Plasticisers are used to make the product they are added to softer, flexible and easily workable; plastic materials acquire a long series of peculiarities when the plasticisers are added [1]. The definition of plasticisers, established by the Council of the International Union of Pure and Applied Chemistry (IUPAC), is: *“A plasticizer is a substance or material incorporated in a material (usually a plastic or elastomer) to increase its flexibility, workability, or distensibility”*[2]. Many plastic materials need to be easily worked, flexible and soft for their applications but at the same time resistant to mechanical stress without breaking and able to take different shapes for various needs [3]. Depending on the desired effects and characteristics it is possible to add a small or big quantity of plasticiser; a small addition of plasticiser usually improves the workability of the plastic material while a large addition can completely alter the material properties and its possible applications [4]. A particular aspect is that the plasticisers, once added to a plastic material, are able to lower their glass transition temperature (Tg) [4], in other words the temperature at which the structure of polymer changes from glass state, particularly rigid and fragile, to rubbery state characterized by high flexibility and softness [5]. This temperature depends on the structure and chemical composition of the polymer and can vary enormously among the various classes of compounds; the Tg depends on the macromolecular mobility of the polymer chains, therefore an increased mobility of the chain will be reflected in an increased flexibility and vice versa [6–7]. The polymer with the best compatibility with the plasticisers is *poly-(vinyl chloride)* or PVC, this can be used even without the addition of plasticisers as unplasticised PVC (PVC-U) for the production of pipes and siding. The PVC with the addition of plasticisers can be used in many fields such as coating of electrical cables, flooring, automotive interior trim, clothing, covers, paints, food packaging, fixtures, medical materials and much more [4]. Over 90% of plasticisers produced in Europe are used for the production of plasticised PVC to create a whole range of flexible and durable applications [3].

### 1.2 General plasticisers

Plasticisers can be essentially divided into monomeric plasticisers, in general low molecular weight compounds, and polymeric plasticisers, obtained by low molecular weight monomer polymerisation [1]. The monomeric plasticisers are divided into six families depending on the leader compound they are related to (Fig.01): phthalate (C_8_), adipate (C_6_), trimellitate (C_9_), sebacate (C_10_), azelate (C_9_) and phosphate (PO_4_^3-^). The acids are usually esterified with different alcohols, changing the properties of PVC due to the different interactions between the polymer chains and the plasticiser itself. Currently ~300 monomeric plasticisers are produced but only 100 of them have an economic relevance [4]. In addition, plasticisers can be characterized by their concentration. The primary plasticisers are those used as the only component or as main plasticiser when mixed with others, causing an increasing of elastic behaviours and softness; secondary plasticisers do not lead to significant changes, they are used in smaller concentration and very often combined with primary ones to improve specific performance of the plastic materials, such as reduced flammability [1–2]. The best known and most used plasticisers: are *di(2-ethylhexyl) phthalate* (DEHP), *di(2-ethylhexyl) adipate* (DEHA), *tris(2-ethylhexyl) trimellitate* (TOTM), *tris(2-ethylhexyl) phosphate*, *di(2-ethylhexyl) sebacate* (DOS) and *di(2-ethylhexyl) azelate*(DOZ). Of particular interest are the plasticisers obtained from the esterification of *adipic acid* (C_6_) and *primary alcohols* C_7_-C_10_ differently branched. The group of the adipate esters, in relation to phthalates, can show better performance at low temperature [1–4]: they extend the temperature range to which the PVC remains flexible and therefore they drop considerably the Tg of the polymer. The adipate esters are used to produce film suitable for food packaging, coating of electrical cables, soles for shoes and more. The most used are *di(2-ethylhexyl) adipate* (DEHA), *diisononyl adipate* (DINA), *di(2-butoxyethyl) adipate* (DBEA) and *dioctyl adipate* (DOA). They are not usually used as unique plasticiser component of PVC, but they are combined with phthalates in order to have a compromise of imprinted characteristics, but overall they are expensive. As a matter of fact the interest is to reduce the production cost of these molecules and to make the productive process more sustainable, not only for the esters but also for the adipic acid and alcohols used [8].

**Fig. 01.**
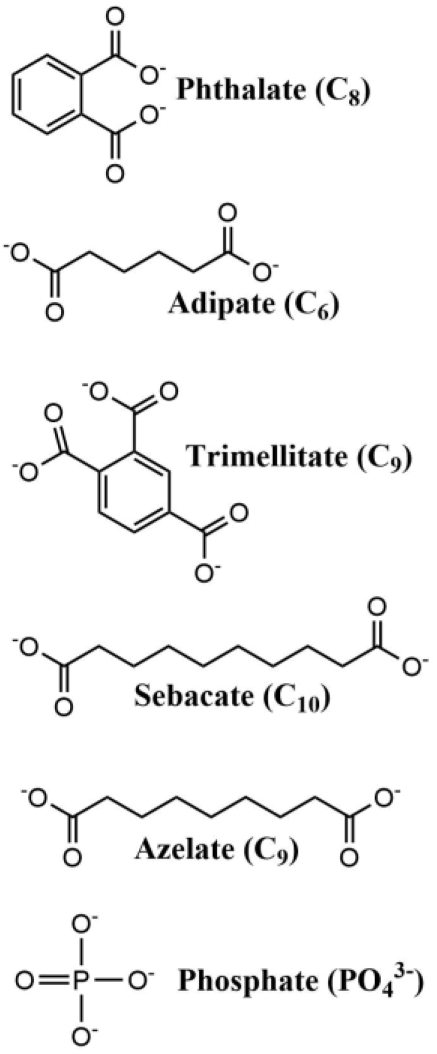
families of the monomeric plasticisers; image *by Andrea Pasquale*.

### 1.3 Usage and commercial relevance of adipate

About 7.5 million tons of plasticisers are consumed worldwide every year and Europe consumes more than 1.3 million tons [9]. Adipic acid, a linear dicarboxylic acid composed of six atoms of carbon, is considered as the major dicarboxylic acid on the market and it is a fundamental chemical building block for the synthesis of lots of molecules [10]. It is classified as a medium-chain dicarboxylic acids (MDCAs) with high added value [11]. About the 65% of the adipic acid produced worldwide is used to produce *Nylon-6,6-polyamide* and most of it derives from fossil sources [12]. Adipic acid is also used to produce polyurethanes, paints, coatings, plasticisers and also as food additive to give an acid taste (E355) [13]. The global market for adipic acid was 2,610 kilo-tons in 2012 and it is expected to reach about 3,747.2 kilo-tons by 2020, growing 4.7% from 2014 to 2020. In terms of revenue, the market for adipic acid was valued USD 4,315.6 million in 2012 and it is expected to reach USD 7,240.8 million by 2020, growing of 6.2% from 2014 to 2020 [8]. Pacific Asia drove the global market of adipic acid for about 35% of the total volume in 2012. Europe is the second major producer worldwide of adipic acid for about the 27% in 2012.

### 1.4 Chemical synthesis of adipic acid

The most widely used method to produce adipic acid is chemical synthesis from fossil-based chemicals at industrial level. It consists in the reduction of benzene followed by two steps of oxidation [14], but adipic acid is also made by the hydrocyanation of butadiene, followed by hydroisomerization in adiponitrile, which is then hydrolysed to adipate [13]. The synthesis reaction (scheme 01) starting from benzene (OHr,) has a first step of reduction to cyclohexane (C_6_H_12_) at the optimal reaction conditions (Ni-Al_2_O_3_, H_2_, 2600-5500 kPa). Subsequently cyclohexane is oxidized to KA-Oil (ketone-alcohol oil) or cyclohexanone (C_6_H_10_O) and cyclohexanol (C_6_H_11_OH) in the presence of Co, O_2_, 830-960 kPa e150-160°C. Finally, the KA-Oil reacts with nitric acid in air producing adipic acid through a degradative oxidation (Cu, NH_4_VO_3_, 60-80°C) [15]; this last step of oxidation is the cause for the release of nitrous oxide (N_2_O) as a by-product of reaction and a whole series of greenhouse gas (GHG) such as nitrogen dioxide (NO_2_) and nitrogen monoxide (NO). The chemical oxidation reactions, as shown, are very polluting, also the use of heavy metals such as Cu and Co can be dangerous, they need appropriate disposal.

**Reaction scheme 01.**
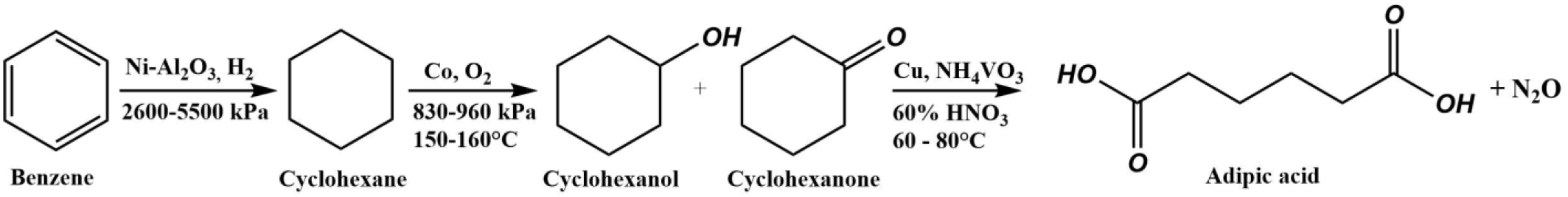
chemical synthesis of adipic acid by reduction of benzene followed by two steps of oxidation; image *by Andrea Pasquale*.

### 1.5 Sustainability of adipic acid for the environment

Nitrous oxide (N_2_O) is nearly 300 times more polluting than carbon dioxide (CO_2_) [10]. The production of adipic acid is responsible for the emission of some quantity of nitrous oxide N_2_O in atmosphere that turns out to be an inevitable stoichiometric waste of the reaction and it is the cause for greenhouse effect and ozone depletion. Despite the best technologies for the recovery of nitrous oxide, about 400.000 metric tons are released every year by the production [17], that matches about 5-8% of the total anthropogenic emission in the world [16]. The conditions of the reaction are not mild but require high temperature and pressure to develop, furthermore multiple reaction steps are expected, not a single step; for all these reasons, in addition to emission, the process can be considered unsustainable and highly polluting. Moreover, the production process presents a high complexity, high energy consumption and release of by-product [18]. Because of the growing awareness for the environmental protection and for the creation of a circular economy, the aim is to develop a sustainable synthesis process, also focusing on bio-based processes centered on the reuse of waste material. The compounds composed of two to six atoms of carbon, with different function, are considered as very valuable chemical compounds deriving from biomass, accessible through fermentation and biotransformation [19]. From the analysis of the economic opportunities for the production from glucose, the adipic acid results to be an interesting compound for a bio-based process [18]. The development of a bio-based process for the production of adipic acid from waste materials would reduce the energetic costs of production, the carbon footprint compared to oil, the emissions of GHG and decrease of CO_2_. Such a bio-based process would allow for a large metric reduction for life cycle assessment (LCA) related to human health, quality of ecosystem and depletion of resources [20].

## 2. Biosynthesis of adipic acid

Plasticisers can be used to resolve the problems that are found in biopolymers, and they are now considered as a promising materials for the future [21]. Plasticisers reduce the stress-strain, hardness, viscosity and the creation of electrostatic charges on the polymer, secondly, they increase the flexibility of chains, fracture toughness and the dielectric constant [22].

### 2.1 Direct oxidation of cyclohexenes

An alternative to the chemical oxidation of adipic acid is the direct oxidation of cyclohexene with hydrogen peroxide (H_2_O_2_) at 30%, as reported by *R. Noyori et al.*[16]. The hydrogen peroxide is an excellent and clean oxidant in the oxidation reactions, when it is used at concentrations lower than 60%. The described reaction consists of a mixture of: cyclohexene (C_6_H_10_), H_2_O_2_, sodium tungstate Na_2_WO_4_*2H_2_O and methyltrioctylammonium hydrogen sulphate [CH_3_(n-C_8_H_17_)_3_N]HSO_4_ as a phase-transfer catalyst (PTC); the mixture was stirred in air at 1000 rpm and at a temperature of 98°C for 8 hours. This reaction permits to obtain adipic acid with a yield of 93%, after filtration and drying it get an analytically pure yield of 90%. Now companies are still working for developing an efficient process to produce hydrogen peroxide in order to be used as the clean oxidant [23]. With this method is possible to have a directed reaction but many different components are necessary, as the sodium tungstate (Na_2_WO_4_*2H_2_O) and methyl-trioctylammonium hydrogen sulphate ([CH_3_(n-C_8_H_17_)_3_N]HSO_4_).

### 2.2 Biosynthesis bio-based

Between the various possible solutions for the sustainable production of adipic acid, there is the bio-based process using waste materials as substrate. Bio-based plasticisers have several advantages, such as renewability, degradability, hypotoxicity and excellent solvent resistance extraction, making them potential to replace *o*-phthalate partially or totally [24]. The ultimate goal is to obtain a consolidated bioprocess (CBP), taking advantage of the ability of some organisms to directly synthesize specific molecules [25–26]. These processes require the use of specific microorganisms capable of tolerate the industrial conditions, fulfilling the yield, production and productivity parameters. *E. coli*, *S. cerevisiae* and *C. glutammicum* were investigated for this purpose, because they are the most used biological systems, known at omic-level and easy to engineer [10]. Adipic acid’s pKa is 4.4, so when it is released in the culture broth the pH decreases. Therefore, the microorganism must be able to tolerate high concentration of acids. For this, it seems that *S. cerevisiae* would be the most suitable microorganism, because it can maintain high yield even in acid environment during the production of ethanol [27]; furthermore, yeast is an interesting microorganism for metabolic engineering in order to produce adipic acid. *E. coli* is able to survive in acid environment, but is very sensitive to pH variations, therefore it may not be adapt for this process. The same reasons can apply for *C. glutammicum*, which is normally cultivated at neutral pH [28]. At the moment no microorganisms able to naturally produce adipic acid are used, but a bacterium that presents a possible pathway for adipic acid synthesis, *Thermobifida fusca*, has recently been discovered [29]. In order to maintain the high yield and the low costs of production, choosing the most appropriate microorganism is of vital importance. Many metabolic strategies have been adopted in order to directly produce adipic acid or its precursors which are then processed, through direct or indirect fermentations.

The first method was to produce adipic acid via indirect fermentation of *cis,cis*-muconic acid or glucaric acid, later converted in adipic acid in a hydrogenation process using Pt as catalyst [13]. *Cis,cis*-muconic acid is a dicarboxylic acid with two conjugated double bonds which is an intermediate of the degradation pathway of benzoate (KEGG map00362), and it is relevant for the chemical industry [30]. It can be obtained via glucose or benzoate fermentation: the β-ketoadipate pathway, a widely used degradation pathway for aromatic molecules such as benzoate, is relevant for the synthesis of *cis,cis*-muconic acid. In this pathway benzoate is converted to catechol (C_6_H_6_O_2_) following the ortho-cleavage pathway, then converted to *cis,cis*-muconic acid and in the end to succinyl-CoA and acetyl-CoA [31]. Starting instead from glucose, an artificial biosynthetic pathway was created, in which the 3-dehydroshikimik acid (DHS), an intermediary of the biosynthesis of aromatic amino acids, is converted into *cis,cis*-muconic acid using a strain of *E. coli* lacking the *shikimate dehydrogenase enzyme*, fundamental for amino acid biosynthesis [32]. Concerning the production of glucaric acid, the co-expression in *E. coli* of genes coding, in order, for myo-inositol-1-phosphatesynthase *(Ino1)* from *S. Cerevisiae*, for mio-inositol oxygenase (*MIOX*) from mice, and for urinate dehydrogenase (*udh*) from *P. Syringae*, allows the accumulation of the acid, which is subsequently subjected to chemo catalytic hydrogenation to obtain adipic acid [33].

Other works have focused on the engineering of a strain of *S. Cerevisiae* so that it may produce a recombinant enzyme, an enoate reductase (*ERs*), capable of performing the hydrogenation enzymatic reaction of the *cis,cis*-muconic acid once it’s been synthesized by the same microorganism. *S. Cerevisiae* was engineered with three plasmids in order to create a biosynthetic pathway for the production of adipic acid; the fermentation consists of three steps: the growth of the cells, the production of the enoate reductase (*ERBC* enoate reductase from *Bacillus coagulans)* along with the biosynthesis of the muconic acid, and lastly the production of the adipic acid [34]. The production yield is not very high, 2.59 ± 0.5 mg/L, but that of the *cis,cis*-muconic acid is much higher, indicating the possibility to improve yield and enzymatic efficiency.

Another attempt to produce adipic acid exploits the combination of the direct or inverse β-oxidation with the ω-oxidation [35]. Such trials have been made with *Candida spp*. and *E. coli*. In the case of *Candida spp,* in order to combine the two pathways (ω-oxidation and β-oxidation), a key issue is the selection of a specific acyl-CoA oxidase (*POX5*) to be involved in the first reaction of the β-oxidation, as this enzyme doesn’t normally accept dicarboxylic acids. During the processing of the palmitic acid (C16), it is converted in hexadecanedioic acid through ω-oxidation, then the β-oxidation with *POX5* produces adipic acid, secreted in the growth medium [36]. The achieved titer of adipic acid is 50 g/L [10], quite high with respect to the other methods. Another trial takes advantage of the implementation of the 2-oxopimelic acid pathway in *E. coli*, along with the alternative path of synthesis of lysine passing by the 6-aminocaproic acid as an intermediary. Both paths bring to the formation of the adipic acid semialdehyde, which, once oxidized, generates the adipic acid as a by-product [37]. The pathway allows to obtain adipic acid at the relatively low titer of 0.3 g/L; furthermore, in the production of one mole of adipic acid, starting from 2-oxoglutarate, one mole of NADH is consumed, resulting in an imbalance of the redox potential of the reaction. This may be a reason why very little of the product of interest is synthesized.

Other works describe also the method by which the succinyl-CoA can be converted into adipic acid through the system of polyketide synthase (PKS) [38]. In this case the synthesis hasbeen performed with isolated enzymes using the succinyl-CoA as the starting molecule, and the malonyl-CoA as carbon additive unity. This path has been chosen for being capable of extending the succinyl-CoA to a 3-hydroxy-adipyl-ACP, which can be easily converted to adipic acid with the use of three enzymes [39]. Also in this case the yield is low, approximately 0.3 g/L, and for each molecule of acid produced two equivalents of NADPH are consumed, causing an imbalance in the redox potential [10].

### 2.3 Reverse adipate degradation pathway RADP

Among all the metabolic strategies that have been described, the reverse adipate degradation pathway (RADP) is the most performing, it can produce adipic acid at the titer of 68 g/L by fed-batch direct fermentation and using glucose as carbons and energy resource, with a yield of 93.1%. This path was identified for the first time in 2015 in *Thermobifida fusca* by *Niti Vanee et al.*, during the characterization of the microorganism’s proteome and metabolome [40]. The pathway is composed of five reactions, catalyzed by five specific enzymes: *Tfu_0875* (β-ketothiolase), *Tfu_2399* (3-hydroxyacyl-CoA dehydrogenase), *Tfu_0067* (3-hydroxyadipyl-CoA dehydrogenase), *Tfu_1647* (5-carboxy-2-pentenoyl-CoA reductase) and *Tfu_2576-7* (adipyl-CoA synthetase). The pathway uses as feedstocks acetyl-CoA, derived from the complex of pyruvate dehydrogenase, and succinyl-CoA, as the metabolic intermediate of the Krebs Cycle; with the succession of the reactions, using as cofactors NADH and FADH_2_, the direct production of adipic acid at high yield is possible, using glucose as resource of carbon and energy, that can derive from waste materials and renewable resources (figure 02). The work by *Mei Zhao et al.* allowed to use *E. coli* as system for the productions of adipic acid, by making some modifications to the strain [41]. It was necessary to over express the enzyme *Tfu_1647* because it was identified as the bottleneck that limits the final production of adipic acid, passing by a yield of 11% to 49.5%, with only this change. Furthermore, by using CRISPR/Cas9 the major pathways of flux carbon have been eliminated/limited, versus the synthesis of adipate: the gene *ldhA* of L-lactate dehydrogenase was deleted to remove the production of acid lactic, the gene *sucD* encoding for succinyl-CoA synthetase was deleted to promote the accumulation of succinyl-CoA and the gene *atoB* encoding for acetyl-CoA C-acetyltransferase, involved in the reaction for produce acetoacetyl-CoA, was deleted. The *E. coli* BL21 (DE3) strain used is *ΔldhA, ΔatoB, e ΔsucD* and engineered with pUC57 plasmid built to contain all the coding sequences of the enzymes of the reverse adipate degradation pathway [41]. The production of adipic acid by *E. coli* in a 5L bioreactor was conducted at 37°C for culturing cells and at 30°C for inducing gene expression with 1 vvm aeration and 400 rpm stirring with 1mM IPTG. From various experiments, adipic acid was obtained with a maximum titer of 68 g/L after 88h on fed-batch fermentation, with OD600 maximum of 36.7 after 64h. Finally, using a specific method with eight steps, the purification of adipic acid was possible and it could be solubilized in water for the analysis at mass spectrometry.

**Fig. 02.**
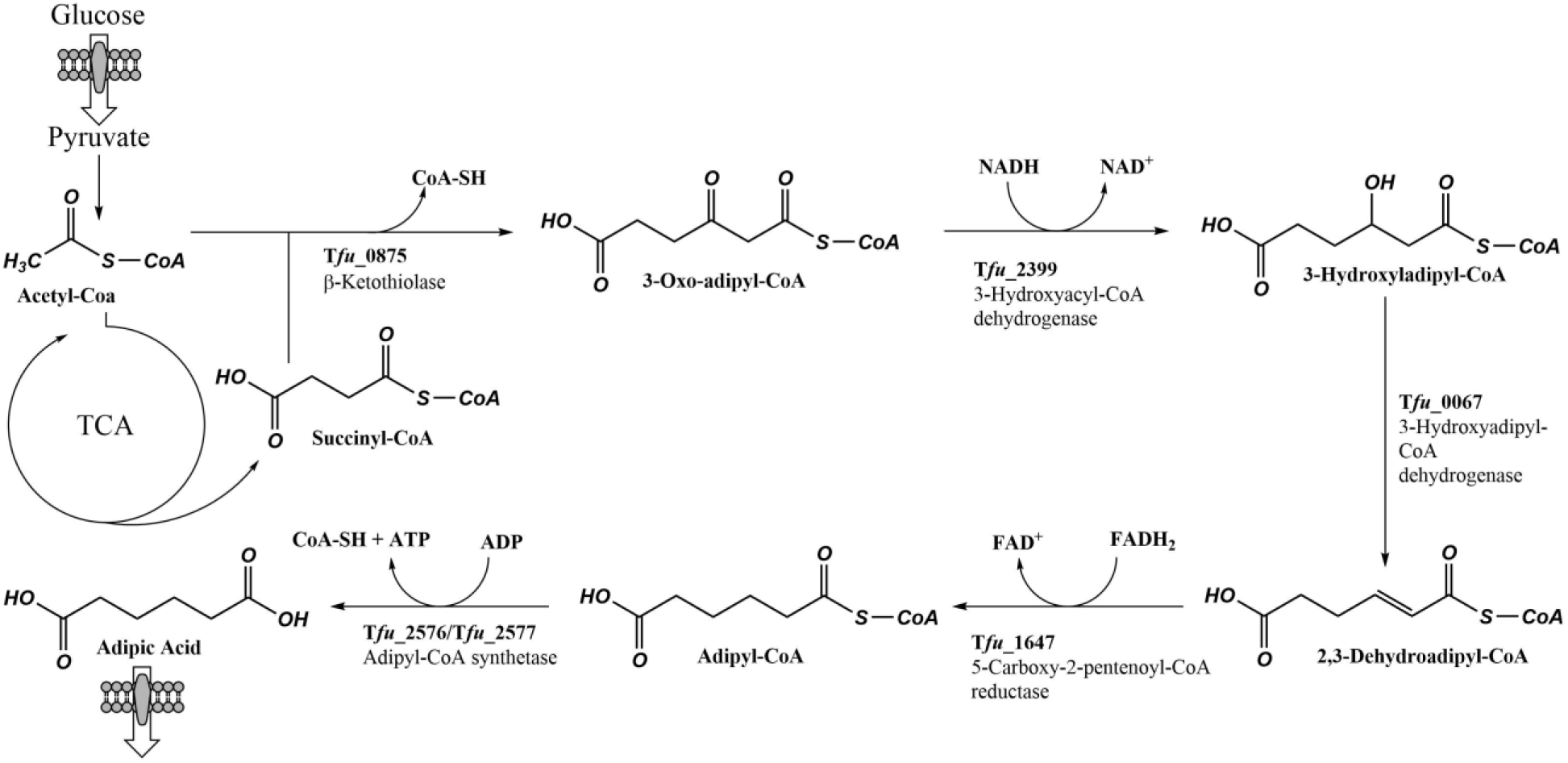
metabolic pathway. the reverse adipate degradation pathway (RADP) includes five steps in *E. coli*. The pathway uses feedstocks acetyl-CoA, derived from the complex of pyruvate dehydrogenase, and succinyl-CoA, as the metabolic intermediate of the Krebs Cycle. At the end adipic acid is released into extracellular environment: image *by Andrea Pasquale*.

## 3. Biocatalytic reactions to produce adipate esters

### 3.1 Properties of DEHA and chemical synthesis

Di(2-ethylhexyl) adipate (DEHA) is one of the most used adipic acid ester, as it guarantees good oxidation stability, low toxicity, low volatility, high viscosity index and high biodegradability. Furthermore, it has been used in many applications such as paint stripper, fragrance, lubricant, food packaging and plasticiser [42]. In particular, using it as a plasticiser provides various unique properties for PVC, such as flexibility, elasticity and workability; in addition, it is less sensitive to temperature changes, as it is more fluidic at low temperatures and less volatile at high temperatures [43]. Another plus is that it is more ecofriendly when compared with phthalates.

Chemical synthesis of this molecule is carried on by Fischer esterification of adipic acid and monohydric alcohols, using corrosive chemical catalysts as methanesulfonic acid, cation-exchange resins and modified heteropoly acids [42], under high reaction temperature for which complex and expensive facilities are required. In addition, it is necessary to use a higher quantity of raw materials because Fischer esterification is an equilibriumreaction where reagents are not fully converted to products, resulting in higher costs and high waste generation. Considering this disadvantages, biocatalytic processes could offer significant advantages because of milder reaction conditions, lower energy requirement, the ease of product isolation and the possibility to reuse the catalyst [44].

### 3.2 *Candida antarctica* lipase-catalysed synthesis

Lipases (EC 3.1.1.3: Triacylglycerol acyl hydrolase) constitute the third most important category of enzyme and their global market size is projected to reach $590.5 million by 2020 [45]. They can be used to catalyze fatty-acid esters synthesis in nonaqueous solvent, for which lipases are frequently tolerant. Therefore, they can be used as biocatalyst for DEHA synthesis from one mole of adipic acid and two mole of 2-ethylhexanol (scheme 02).

**Reaction scheme 02.**
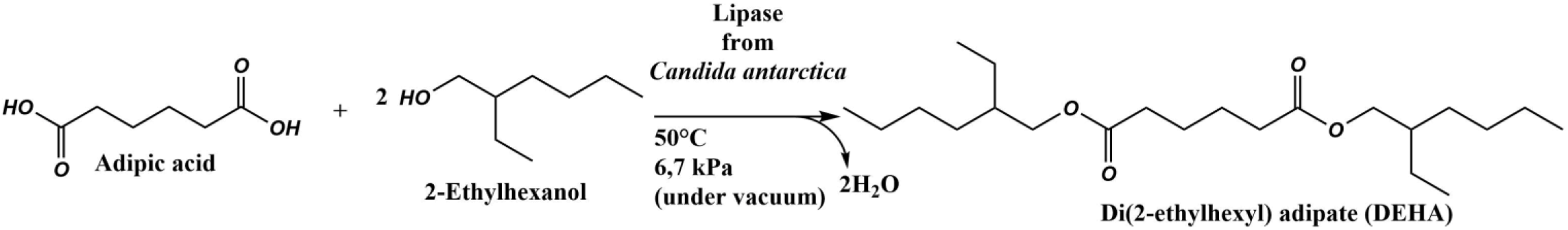
biocatalytic reaction to produce adipate esters DEHA by lipase from *Candida antarctica*; images *by Andrea Pasquale*.

A lipase from *Candida antarctica* can be used. Its optimal reaction conditions were investigated to obtain the maximum yield, production and productivity for the process.

The conversion to DEHA in the reaction of adipic acid with 2-ethylhexanol (mol%) was calculated as follows:

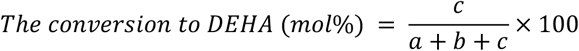

Where *a* represents the moles of adipic acid in the reaction mixture, *b* the moles of mono ethylhexyl adipate (MEHA) in the reaction mixture and *c* the moles of DEHA in the reaction mixture [46], therefore it is important to consider non-reacted substrates and byproducts, decreasing the total yield.

### 3.3 Lipase characterization

*Candida antarctica* lipase was biochemically characterized. In several studies, where esterification of adipic acid and various alcohols was carried out, it was evident that an increase in temperature between the given range (35-65°C) improved solubility of the substrates and reduced viscosity, which enhanced the yield of conversion to DEHA [47]. However, increasing the temperature may reduce the stability of an enzyme and would prevent its full exploitation. Furthermore, the energy requirement and the costs increase to maintain the higher reaction temperature. Thus, after stability tests, 50°C was selected as the optimal temperature [46].

Enzyme loading is another crucial factor to consider: in general, a large amount of enzyme increases the reaction rate as well as the yield, but it increases the cost, so trade-offs must be evaluated. Thus, 5% was selected as the optimal enzyme loading [46].

Lastly, reuse of the enzymes was investigated by evaluating their residual activity, which began to decrease steadily from the 17^th^ cycle, due to various factors, such as shear stress and high temperature that leads to the denaturation of the lipase. After 33 cycles, the residual activity decreased by ca. 81% of its original activity [46], showing a good stability of the lipase.

### 3.4 Immobilized enzyme

When one mole of adipic acid reacts with two moles of 2-ethylhexanol, two moles of water are produced along with one mole of DEHA. The presence of water can be undesirable for the process, because it can cause lipase-catalyzed hydrolysis if the water formed from the esterification is not removed effectively from the reaction system, even though a minimum amount is required to maintain the lipase structure for the catalytic activity [48].

In order to guarantee these conditions, a solvent-free system under vacuum condition, where enzymes are immobilized, is applied. This system brings various advantages, such as: increasing the recovery yield of the final product, which is defined as the mass of product formed per unit of reactor volume per hour, that reduces the cost of the bioreactor designed for large-scale productions; minimizing environmental impact by avoiding the use of toxic and flammable organic solvent and, lastly, significantly reducing the costs due to fewer purification steps in downstream processes [42].

For production of DEHA in specific, the conversion to DEHA reached 100 mol% under vacuum condition within 4h. In contrast, without vacuum maximum conversion to DEHA was only 75 mol% after 9h [46]. Up to a certain level of vacuum, reaction rate increases because water is removed efficiently, however, excessive vacuum causes a decrease of reaction rate, because lipase-catalyzed reactions require essential water for catalytic activity of lipase [49], as a result 6.7 kPa was selected as the optimal pressure condition [46].

Mixing and mass transfer in the reaction system are factors to investigate, considering the high viscosity of the mixture in the absence of solvent. The results showed that employing above 100 rpm agitation speed with a stirrer was necessary [42].

It is important to consider the stoichiometry of the reaction: the molar ratio of 2-ethylhexanol and adipic acid must be 2:1 to achieve a maximum yield of DEHA [46]. However, a high molar ratio of 2-ethylhexanol has been suggested to be a requirement as it may improve the dispersion of the adipic acid in the reaction mixture, especially in the case of the solvent-free system [50]. Hence, a molar ratio of 2.5:1 was considered to be optimal [46], because in this condition the reaction is complete and monoethylhexyl adipate (MEHA) does not form. 2-ethylhexanol is the only non-reacted byproduct and its removal is more energy-efficient, since a single step of distillation is sufficient and its boiling point is lower than that of MEHA, the conversion to DEHA reached 100 mol% after this only one downstream step.

## 4. Conclusion

In summary, among all the analyzed pathway for the production of adipic acid, the Reverse Adipate Degradation Pathway (RAPD) was the most performing one and the one with the highest yield of 93.1%. *E. coli* was selected as the cell factory, but using *S. cerevisiae* could be an interesting choice for its tolerance towards low pH. In comparison to the chemical synthesis, direct fermentation is more sustainable, because: it releases lower N_2_O and other greenhouse gasses in the atmosphere, heavy metals are not required, reaction conditions are mild and fewer reaction steps are required. One of the future aim is to reduce the downstream steps for recovery of adipic acid and improve the technologies in order to lower the cost of the process. In order to scale-up the process for industrial production, glucose could be derived from second generation feedstocks, such as waste materials from agriculture or organic waste.

For the production of a sustainable plasticiser, the second step is the esterification of one mole of adipic acid with two moles of 2-hethylhexanol by a lipase from *Candida antarctica*, obtaining di(2-ethylhexyl) adipate (DEHA), one of the most used plasticiser for PVC. Using a solvent-free system with immobilized enzymes the conversion to DEHA reaches 100 mol%, and only one purification step is required for the downstream. Applying reduced pressure to the reaction increased its yield, therefore it could be used in other enzymatic reaction to make them more efficient and sustainable. For the purpose of reducing the reaction time (a key issue in industrial production), protein engineering could be a solution to improve the catalytic activity of the enzyme; an alternative solution is bio-prospecting for enzymes produced by other microorganism with higher capacities, for example enzyme from psychrophilic microorganism in order to lower the reaction temperature and save energy costs.

The perspective is to develop an industrial process combining the above mentioned process for adipic acid synthesis and esterification in order to synthesize DEHA with high productivity, starting from waste materials. These results demonstrate that biotechnology is relevant for the plasticisers industry for developing a more green and eco-friendly production process.

